# Uncovering Viral Signals Across Cell Populations in Colorectal Cancer

**DOI:** 10.64898/2026.11.07.765073

**Authors:** Mohammadamin Mahmanzar, Karim Rahimian, Kaveh Kavousi, Youping Deng

**Affiliations:** Department of Quantitative Health Sciences, John A. Burns School of Medicine, University of Hawaii at Manoa, Honolulu, HI 96813, USA; Department of Bioinformatics, Kish International Campus, University of Tehran, Kish, Iran; Laboratory of Complex Biological Systems & Bioinformatics (CBB), Institute of Biochemistry and Biophysics, University of Tehran, Tehran, Iran

**Keywords:** Colorectal cancer, Tumour microenvironment, Virus–host interactions, Virome

## Abstract

Viral sequences have been detected in colorectal cancer, but their cellular and anatomical distribution remains poorly resolved. We aimed to establish a cell-type-resolved map of viral reference-associated RNA signals across colorectal cancer, tumour-adjacent normal tissue, and healthy colon.

We repurposed human-unmapped reads from single-cell RNA sequencing libraries generated in ten publicly available studies. We retained cell-barcode and unique molecular identifier information, removed duplicate molecules, and aligned reads to a curated human-associated viral reference using three aligners. Viral signals were analysed across cell types, tissue conditions, and colorectal segments. We used participant-grouped repeated cross-validation to evaluate tissue-condition classification and screened an independent paired colorectal cancer proteomics cohort for compatible peptide matches.

The study included 547 libraries from 112 participants, including 49 with matched colorectal cancer and tumour-adjacent normal samples, and approximately 415,000 cells spanning ten major cell populations. Viral reference-associated RNA signals differed across epithelial, immune, stromal, and vascular compartments, tissue conditions, and anatomical regions of the colon. Of 318 independent healthy-reference viral-load comparisons, seven involving six virus– cell-type groups remained significant after Benjamini–Hochberg correction, including plasma-cell HHV-7 and HHV-6B and mast-cell HCV-2. Cell-type-resolved viral features distinguished colorectal cancer, tumour-adjacent normal tissue, and healthy colon in participant-grouped validation; random forest achieved a macro-AUC of 0·957 (SD 0·028), with HERV-K113 as the leading quantitative feature. An independent proteomic screen identified peptide matches compatible with 20 candidate viral groups.

We established a cell-type-resolved map of viral reference-associated RNA signals across colorectal tissue conditions and anatomical regions. By identifying candidate viral signals in epithelial and immune compartments, this map provides a basis for investigating virus–host pathways involved in tumour biology, immune regulation, and antigen-presentation states, and could help prioritise future combination strategies pairing virus-directed or virus-mimetic approaches with standard treatment or immune checkpoint blockade.

## Introduction

Single-cell RNA sequencing (scRNA-seq) has transformed cancer research by resolving gene expression across distinct cell populations ^1^. However, conventional analyses focus on human-mapped reads and generally exclude sequences that fail to align with the human reference genome, although large human RNA-seq studies have reported that approximately 9–20% of reads remain unmapped ^2^. In barcode-based scRNA-seq, recovered unmapped reads can retain a cell barcode and unique molecular identifier (UMI), allowing them to be linked to their cell of origin and examined for additional signals, including nonhuman sequences and endogenous elements ^3^. This creates an opportunity to investigate overlooked molecular features across the epithelial, immune, and stromal compartments in colorectal cancer ^1^.

Viruses can influence host biology through several mechanisms, including changes in RNA stability, integration of viral genetic material into the host genome, and epigenetic regulation ^4–6^. Depending on the virus and infection state, viral genetic material may persist as an episome, integrate into host chromosomes, or be present as an RNA genome ^7^ (Figure 1A). These distinct forms can give rise to viral RNA molecules, including transcripts from episomal or integrated viral DNA, that may be recovered by scRNA-seq when compatible with the library protocol ^8^. However, detecting viral RNAs does not establish genomic integration, episomal persistence, or active replication. Because viral tropism, cellular defenses, and local immunity differ among cell populations ^9^, examining viral read signals at cellular resolution may clarify their distribution across epithelial, immune, and stromal compartments in colorectal tissue.

**Figure 1.**
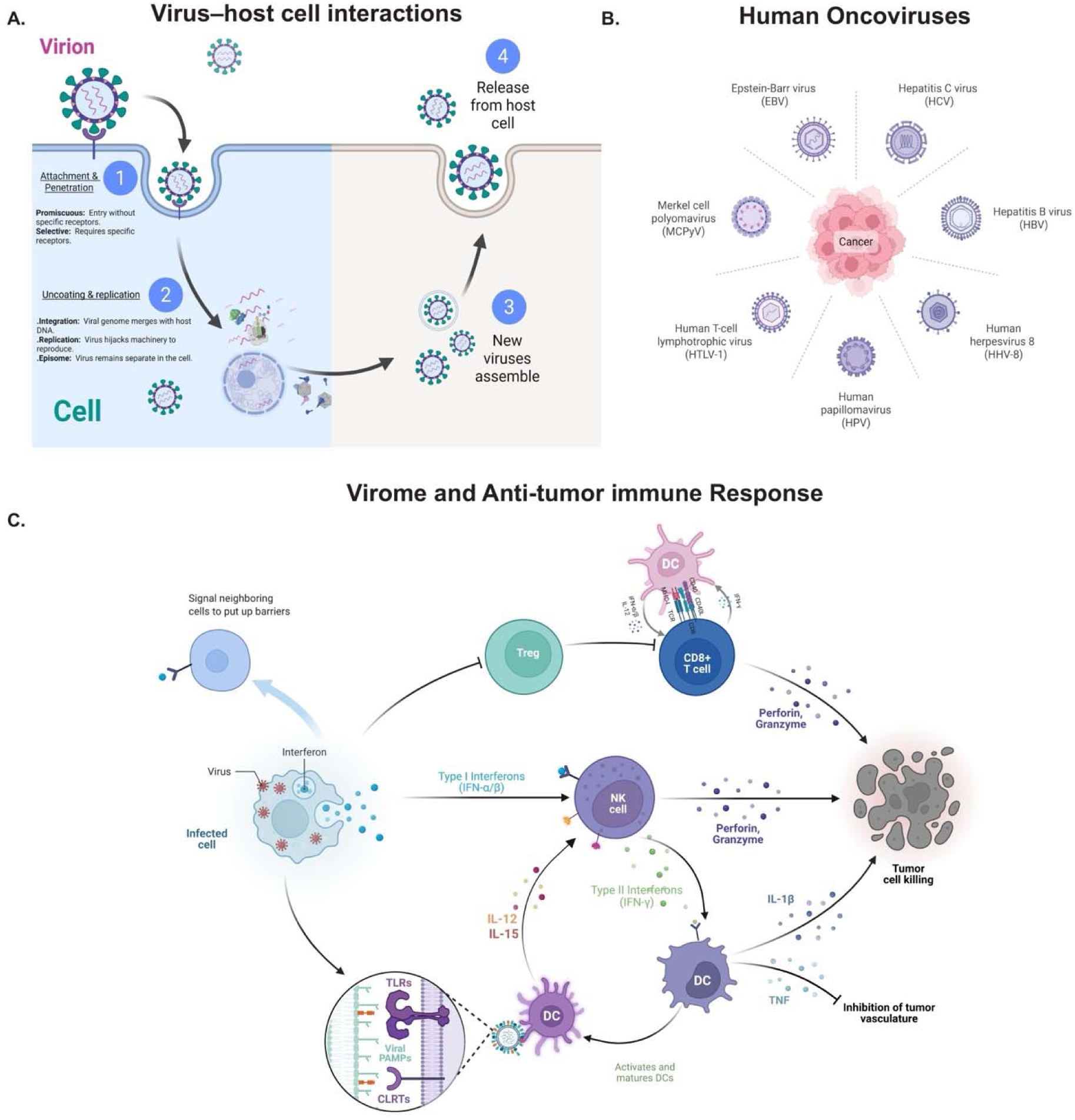
Conceptual framework for virus–host interactions, human oncoviruses and antiviral antitumor immunity. (A) Schematic of the viral life cycle in a host cell, from attachment and penetration through uncoating, genome replication or persistence, virion assembly and release. (B) Examples of human oncoviruses. (C) Conceptual antiviral immune network linking infected-cell sensing and interferon signaling with viral pattern recognition, dendritic-cell activation, natural killer-cell and CD8+ T-cell cytotoxicity, and regulatory T-cell modulation. DC, dendritic cell; TLR, Toll-like receptor; CLR, C-type lectin receptor; PAMP, pathogen-associated molecular pattern; IFN, interferon.

Several viruses are established human carcinogens, including Epstein–Barr virus, hepatitis B and C viruses, high-risk human papillomavirus types, Kaposi sarcoma-associated herpesvirus, human T-cell leukemia virus type 1, and Merkel cell polyomavirus ^5, 10^ (Figure 1B). However, detecting viral footprints in tumours does not establish a direct causal role in cancer but may provide a foundation for further investigation into potential virus–cancer associations. Determining their cellular origins and distributions may help interpret these signals in heterogeneous tissues such as colorectal cancer.

Colorectal cancer (CRC) provides an important setting for investigating viral signals, as previous studies have reported sequences associated with cytomegalovirus (CMV), human papillomavirus (HPV), Epstein–Barr virus (EBV), and the endogenous retroviral locus HERV-K113 in colorectal tumours ^11–13^. However, these sequences have different biological origins. Reads assigned to exogenous viruses may reflect viral material in tissue cells, extracellular RNA, contamination, or ambiguous alignment. In contrast, HERV-K113-associated reads may originate from human endogenous retroviral sequences; HERV-K113 is an insertionally polymorphic locus ^14^. Distinguishing these sources and identifying the cellular compartments carrying the signals are important for interpreting their relevance to CRC.

Virus–host interactions can influence innate and adaptive immune activity in the tumour microenvironment ^15^ (Figure 1C). From a clinical perspective, these interactions may contribute to the distinction between immunologically “hot” tumours, characterized by substantial immune cell infiltration and greater potential responsiveness to immune checkpoint blockade, and “cold” tumours, which exhibit limited immune infiltration and are generally less responsive to immunotherapy. In preclinical colorectal cancer (CRC) models, engineered oncolytic viruses have enhanced T-cell responses and improved antitumour activity in combination with immune checkpoint blockade, including in experimental peritoneal metastasis ^16, 17^. Understanding the cellular distribution of viral signals may provide insights into their associations with the CRC immune microenvironment, including differences between immunologically “hot” and “cold” tumours.

Bulk RNA sequencing combines signals from epithelial, immune, stromal, and vascular cells, making it difficult to determine which populations contribute virus-related reads. Single-cell RNA sequencing could help resolve these signals by linking human-unmapped reads to individual cell populations. However, the cellular and anatomical distribution of viral RNA-associated signals in colorectal cancer remains unclear.

In this study, we aimed to define a cell-type-resolved map of viral signals across colorectal cancer, adjacent normal tissue (ANT), and healthy colon, including paired tumour–ANT samples. We also examined their associations with tissue condition, host-cell states, and colorectal anatomy, and assessed whether selected signals could support future biological and therapeutic investigations.

## Materials and Methods

We developed a barcode-aware workflow to recover viral RNA reads from human-unmapped scRNA-seq data while preserving links among reads, cell barcodes, unique molecular identifiers (UMIs), viral references, and cell-type annotations (Figure 2).

**Figure 2.**
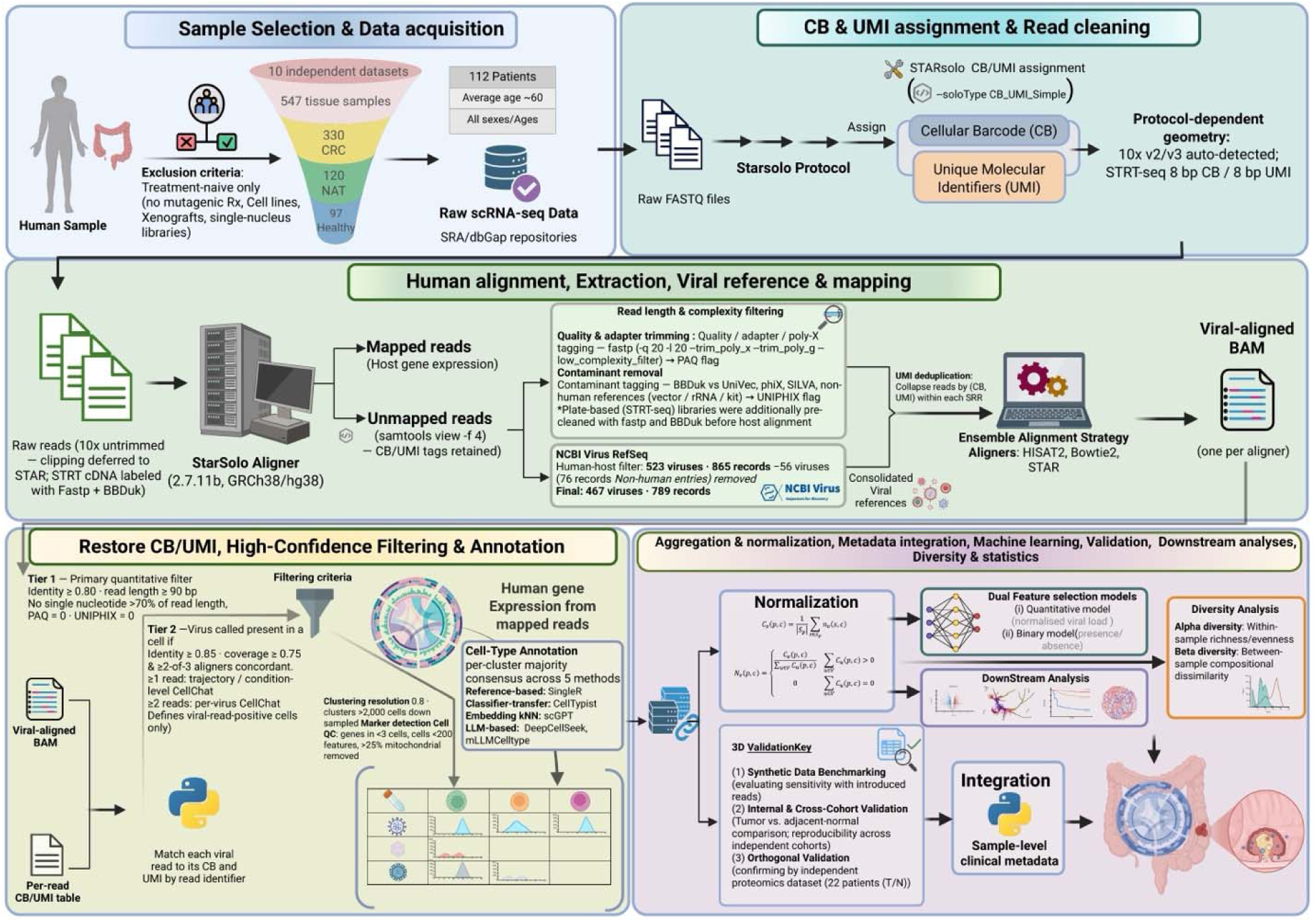
Overview of the single-cell viral-read analysis workflow.

### 1. Study design, data acquisition, and cohort definition

We selected 547 eligible scRNA-seq libraries from ten publicly available studies of colorectal cancer (CRC), ANT, and healthy colorectal tissues, representing 112 biological participants. Cell lines, xenografts, and specimens collected after anticancer treatment were excluded. Following quality control and consensus annotation, the integrated resource contained approximately 415,000 cells. Participant characteristics and library metadata are provided in Tables S1 and S2^18–27^.

Raw FASTQ files were retrieved from the NCBI Sequence Read Archive ^28^ with barcode and technical reads retained as required by each protocol. Of the 547 eligible libraries, 528 were included in analyses requiring viral counts and cell-type assignments (315 CRC, 116 ANT, and 97 healthy).

### 2. Barcode processing, host alignment, and extraction of unmapped reads

Barcode and UMI processing followed the preparation protocol of each library. Droplet libraries were processed using STARsolo ^29^, whereas plate-based libraries underwent protocol-specific barcode processing and read cleaning. Droplet reads were aligned to the Ensembl GRCh38 primary assembly (release 115) using STARsolo to generate Gene and Velocyto features and identify called cells. Reads flagged as unmapped after host alignment were extracted using samtools ^30^ for viral analysis. The primary viral-call filter required a minimum read length of 90 bp 31.

### 3. Viral reference construction, UMI deduplication and viral alignment

A human-associated viral reference ^32^ was constructed from NCBI Virus RefSeq and manually curated, yielding 467 viruses represented by 789 FASTA records (Data S1). Within each sequencing run, reads with valid cell barcodes and UMIs were deduplicated before viral alignment, retaining the read with the highest mean base quality. Deduplicated reads were aligned independently to the curated reference using STAR, HISAT2, ^33^ and Bowtie2 ^34^. Barcode and UMI information was restored after alignment, and evidence was consolidated by sequencing run, read identifier, viral identifier, and cell barcode while recording support from each aligner. Reads mapping to multiple viral accessions were retained for each supported accession.

### 4. Viral call definition and quality filters

Two separate viral-call tracks were used. The primary quantitative track required at least 80% alignment identity, a read length of at least 90 bp, no single nucleotide exceeding 70% of the read, and clear PAQ and UNIPHIX screening flags. This track was used for viral counts, prevalence, diversity, and classification analyses. The stricter cell-positivity track required at least 85% identity, 75% query coverage, and agreement between at least two aligners. It identified positive cells for trajectory and CellChat analyses, requiring at least one supporting read for trajectory and condition-level CellChat analyses and two reads for per-virus CellChat analysis.

### 5. Cell type annotation and attribution of viral reads

Host gene-expression matrices were processed in separate chemistry-aware atlases using Seurat ^35, 36^ for quality control, normalization, clustering, and marker analysis. Cell identities were assigned by consensus across five methods: SingleR ^37^, derived from Human Primary Cell Atlas ^38^, CellTypist ^39^, scGPT ^40^, DeepCellSeek ^41^, and mLLMCelltype ^42^, with discordant annotations reviewed against canonical markers. Fine annotations were consolidated into ten major cell groups. Viral reads were assigned to cell types only when their barcodes directly matched called cells with consensus annotations.

### 6. Metadata integration, paired samples, aggregation and batch structure

Each library was linked to a biological participant, including ^49^ participants with matched CRC and ANT samples. Viral reads were aggregated by library and cell type, then combined across libraries from the same participant. A virus was considered present when at least one retained read was detected in the corresponding cell type. For sample-level analyses, each sequencing run was treated as an observation, accounting for repeated libraries from the same participant. Host transcriptomes were integrated within chemistry-specific atlases. Viral count matrices were batch-corrected using ComBat with tissue condition included in the model. When study and condition were confounded or batches contained fewer than two samples, uncorrected log-transformed matrices were used (Supplementary Methods).

### 7. Downstream host transcriptome and integrative analyses

Differential host-gene expression was analyzed by cell type within separate 10x and STRT-seq atlases. Gene counts were aggregated by sequencing run to generate pseudobulk profiles, and eligible condition comparisons were tested using DESeq2, followed by pathway enrichment with clusterProfiler. All three tissue comparisons were examined in the 10x atlas, whereas STRT-seq supported only CRC versus ANT comparisons. Epithelial trajectories were inferred using Slingshot ^43^, RNA velocity was estimated using scVelo ^44^, and computational spatial projections were generated using Tangram ^45^ and novoSpaRc ^46^. Cell communication was inferred using CellChat ^47^ at the condition and individual-virus levels. Associations between host gene expression and survival were examined using TCGA-COADREAD data.

### 8. Machine-learning analyses

Supervised classification used 189 cell type virus combinations from the primary quantitative track, represented separately as viral load or presence/absence. CatBoost ^48^, random forest, and L2-penalized logistic regression were evaluated using 20 repetitions of stratified fivefold cross-validation. Folds were grouped by biological participant to keep matched CRC and ANT profiles together. Feature construction, scaling, and model fitting were performed within each training fold to prevent information leakage (Supplementary Methods).

### 9. Diversity, viral load and prevalence analyses

Viral diversity and abundance were evaluated using participant-level profiles within each cell type. Alpha diversity was measured using Simpson’s index, with cell-type comparisons performed using Mann–Whitney U tests and Benjamini–Hochberg correction. Beta diversity was evaluated using centered log-ratio (CLR) transformation and Aitchison distance, with differences assessed by PERMANOVA and visualized using UMAP. Viral prevalence was defined by the presence of at least one retained read per virus and cell type, with independent comparisons against healthy tissue evaluated using Fisher’s exact tests. Matched CRC and ANT samples were analyzed separately. Quantitative viral signals were averaged across sequencing runs for each participant and cell type and normalized to the total viral signal within each profile.

### 10. Pipeline recovery, contamination control, and proteomic peptide screening

Pipeline recovery was evaluated by introducing known viral sequences into real sequencing libraries and repeating the workflow (Figure S2). An independent proteomic screen used paired CRC and ANT samples from 22 participants (44 acquisitions; PXD019504). Following the human proteome search, unaccepted spectra or peptides were analyzed with Casanovo and screened against human and microbial or viral protein references. The 1% false discovery rate applied only to the human proteome search.

## Results

### 1. A joint single-cell atlas defines the cellular context of viral signals

We constructed a joint colorectal single-cell t-SNE atlas comprising approximately 415,000 cells from CRC (222,000), ANT (75,000), and healthy tissue (118,000). The atlas resolved ten major cell populations across epithelial, immune, stromal, and vascular compartments, including epithelial, T, B, plasma, macrophage, monocyte/dendritic, mast, neutrophil, stromal/mesenchymal, and endothelial/vascular cells. All three tissue conditions were represented across the major cellular compartments (Figure S3).

We then overlaid viral reference-associated reads onto the atlas to examine their distribution across cell types and tissue conditions. HCV-associated reads were observed across multiple regions of the atlas within three conditions, with genotype 2 showing the most prominent visual signal, alongside genotypes 1 and 6 and isolate H77 (Figure S3). Additional viral candidates, including HPV-71, HHV-6, TBEV, EBV, HSV-1, and Betapolyomavirus macacae, showed varied cellular distributions across tissue conditions (Figure S4).

### 2. Viral read load is structured by cell type and tissue condition

We examined ^12^ selected virus cell-type profiles across CRC (94 participants), ANT (54 participants), and healthy colorectal tissue (13 participants), assessing normalized viral read load, prevalence, demographic distribution, matched CRC–ANT differences, and study contributions (Figures 3A–3G). The complete set of virus–cell-type profiles is provided in Figure S5. Independent comparisons were restricted to CRC versus healthy and ANT versus healthy tissue, whereas CRC and ANT differences were examined among the ^49^ matched participants. All 12 highlighted viral profiles were also detected in ANT tissue, allowing their patterns relative to healthy tissue to be interpreted alongside within-participant differences.

**Figure 3.**
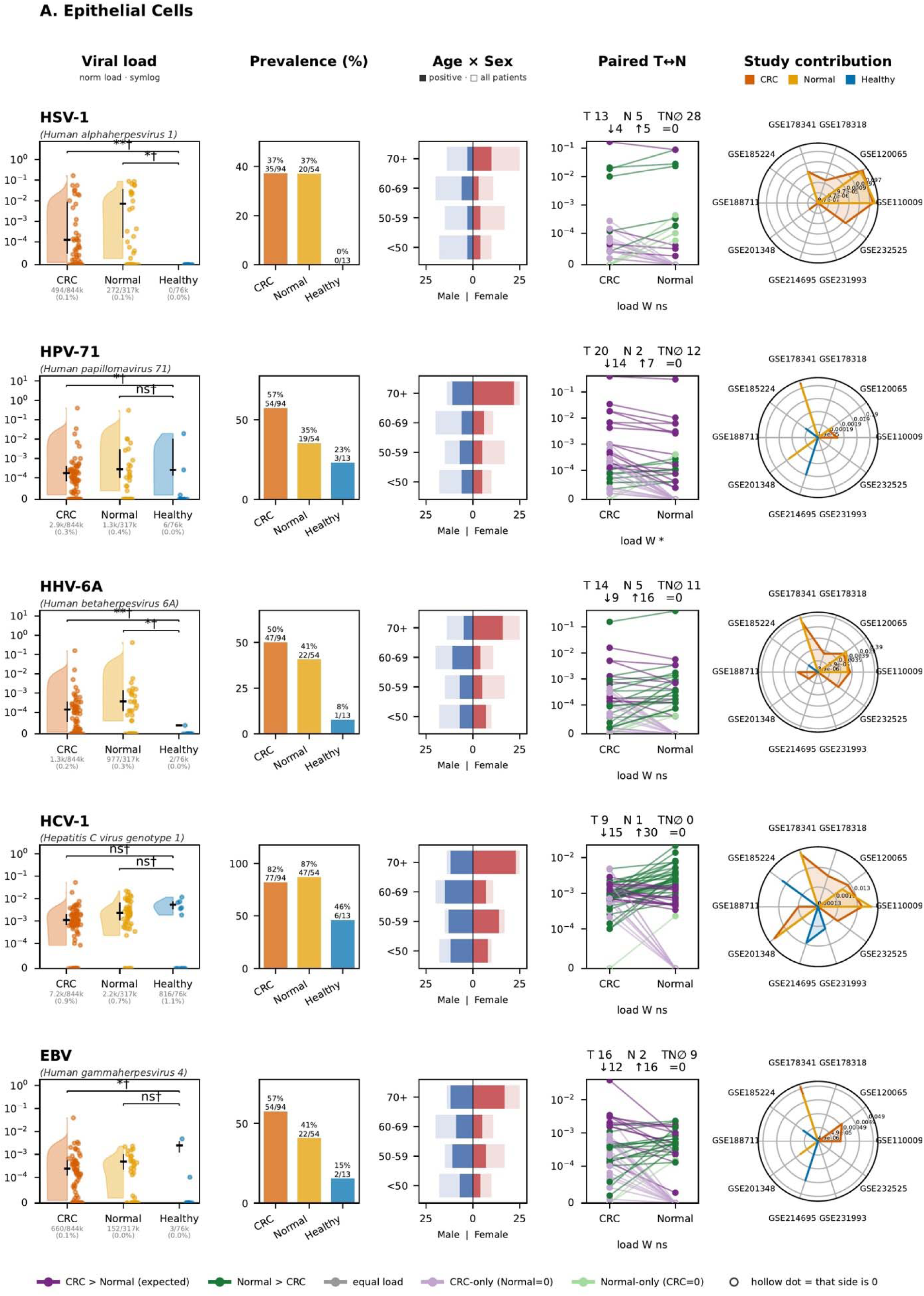
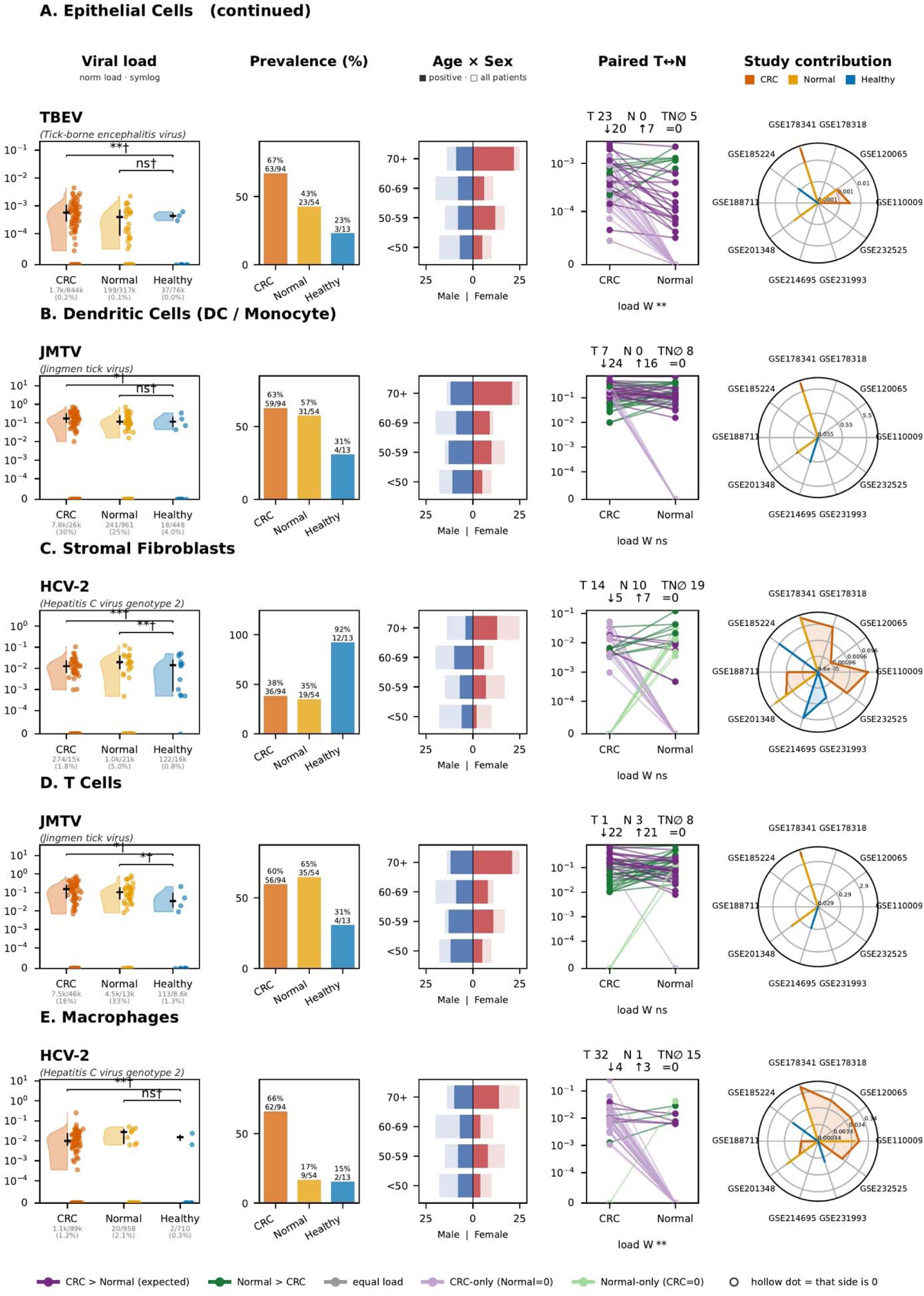
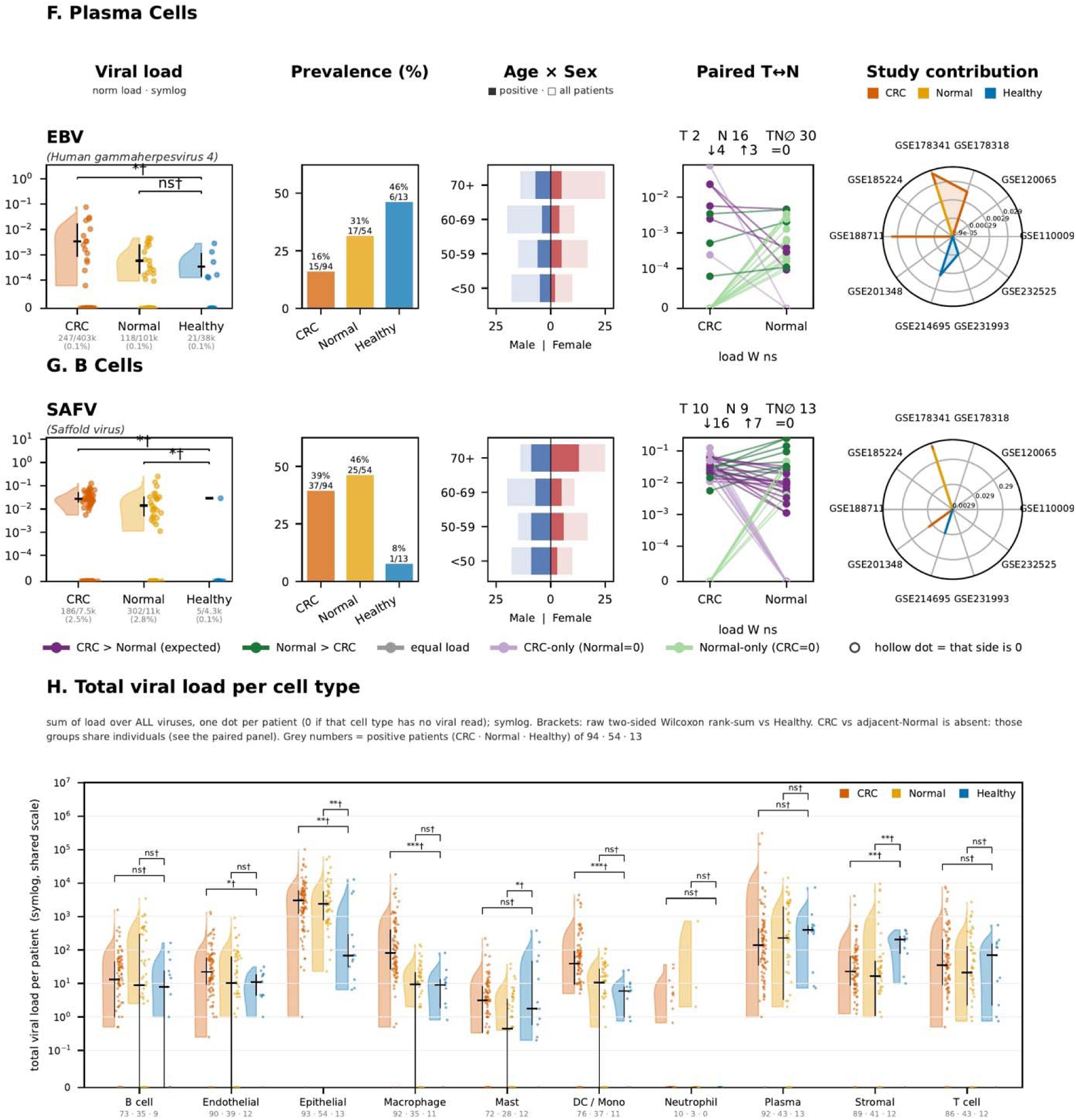
Viral-read profiles across cell types and tissue conditions. **(A) Epithelial cells; (B) monocyte/dendritic cells; (C) stromal/mesenchymal cells; (D) T cells; (E) macrophages; (F) plasma cells; (G) B cells. For (A)–(G)**, each selected virus–cell-type row shows the normalized viral-read-load distribution on a symmetric-log scale, participant-level prevalence, age and recorded-sex distribution, matched CRC–ANT values, and study contribution. Normal in the artwork denotes ANT tissue. Within each displayed virus–cell-type profile, lines connect CRC and ANT values contributed by the same participant. Downward and upward arrows summarize the direction of within-participant change from CRC to ANT, and an equals sign denotes unchanged paired values. W denotes the paired Wilcoxon test; asterisks indicate significance, and ns indicates no significant difference. (H) Summed normalized viral-read load across all viral annotations is shown on a symmetric-log scale for each participant and cell type. Each point represents one participant. \**p <* 0.05; \*\**p <* 0.01; \*\*\**p <* 0.001. Complete virus–cell-type profiles are provided in Figure S5.

#### 2.1 Concordant CRC-associated candidates

Epithelial TBEV and macrophage HCV-2 showed higher viral read loads in CRC than in healthy tissue, with similar patterns in matched CRC–ANT comparisons (Figures 3A and 3E). Epithelial TBEV was detected more often in CRC than in ANT or healthy tissue, and macrophage HCV-2 showed the same CRC-skewed pattern. Both profiles showed higher tumour-associated load in matched comparisons.

Epithelial HPV-71 followed a comparable pattern, with detection in 57% of CRC participants (54/94), 35% of ANT participants (19/54), and 23% of healthy participants (3/13). Viral-read load was higher in CRC than in healthy tissue (*p =* 0.0426), but not between ANT and healthy tissue (*p =* 0.409). The paired analysis also showed a CRC-associated increase (Wilcoxon *p <* 0.05; Figure 3A). Together, epithelial TBEV, macrophage HCV-2, and epithelial HPV-71 showed concordant CRC-skewed patterns across the healthy-reference and matched analyses (Data S2).

#### 2.2 Signals shared between CRC and ANT tissue

Several epithelial viral signals were detected in both CRC and ANT tissue but showed differences relative to healthy colon. HHV-6A and HSV-1 were more common in CRC and ANT than in healthy tissue. Epithelial EBV prevalence was 57%, 41%, and 15%, respectively, with a CRC-versus-healthy load difference (p = 0.0132), but no significant ANT-versus-healthy or paired difference. HCV-1 was highly prevalent in CRC and ANT tissue compared with healthy colon, although viral-read load was similar across conditions (Figure 3A).

Similar patterns occurred in immune compartments. Monocyte/dendritic-cell and T-cell JMTV signals, as well as B-cell Saffold virus, were more common or showed higher load in CRC and/or ANT than in healthy colon, but none showed a significant paired difference. Overall, these profiles showed similarities between CRC and ANT tissue, with the main nominal differences observed relative to healthy colon (Figures 3B, 3D and 3G; Data S2).

#### 2.3 Context-dependent and inverse associations

Stromal HCV-2 and plasma-cell EBV showed contrasting tissue-associated patterns compared with their macrophage and epithelial counterparts. Stromal HCV-2 was more prevalent in healthy colon than in CRC or ANT tissue, with lower viral-read load in CRC and ANT than in healthy tissue, while matched CRC and ANT profiles were similar. Plasma-cell EBV prevalence was also higher in healthy tissue (46%) than in ANT (31%) or CRC (16%) tissue, with a CRC-versus-healthy load difference (p = 0.0308), but no significant differences in the other comparisons. These findings contrast with macrophage HCV-2 and epithelial EBV, highlighting the cell-type dependence of viral-read distributions across tissue conditions (Figures 3C and 3F; Data S2).

#### 2.4 Multiple testing correction across healthy reference comparisons

Among the 318 independent healthy-reference viral-load comparisons, ^63^ were nominally significant (*p <* 0.05), but only seven comparisons involving six virus–cell-type groups remained significant after Benjamini–Hochberg correction (q < 0.05). Four CRC-versus-healthy comparisons remained significant: plasma-cell HHV-7 (q = 0.00115), HHV-6B (q = 0.00190), Punta Toro virus (q = 0.0115), and mast-cell HCV-2 (q = 0.0207). Three ANT-versus-healthy comparisons were significant: plasma-cell HERV-K113 (q = 0.00951), HHV-6B (q = 0.0264), and mast-cell HCV-6 (q = 0.0207). Thus, plasma-cell HHV-6B showed significant differences in both contrasts. In contrast, none of the 24 healthy-reference comparisons for the 12 highlighted profiles in Figures 3A–3G remained significant after correction, with the lowest adjusted *p value* observed for stromal HCV-2 (q = 0.0658). Complete raw and adjusted statistical results are provided in Data S2.

#### 2.5 Aggregate viral load and demographic context

The total viral-read load across all viral annotations showed distinct cell-type and tissue-condition patterns (Figure 3H). Compared with healthy colon, CRC had higher viral-read load in epithelial, macrophage, monocyte/dendritic, and endothelial/vascular cells, and ANT epithelial cells also showed higher load. In contrast, stromal/mesenchymal cells showed higher load in healthy tissue than in CRC or ANT. ANT mast cells also differed from healthy tissue, while the remaining compartments showed no clear differences in the displayed healthy-reference comparisons.

Age and sex distributions showed no consistent demographic concentration across the highlighted viral candidates, although their contributions varied among source studies (Figures 3A–3G). Together, these findings indicate that CRC-associated viral-read patterns differ across cellular compartments rather than reflecting a uniform increase throughout colorectal tissue.

### 3. Viral diversity is structured by cell type and tissue condition

#### 3.1 Participant-level alpha diversity

Participant-level Simpson diversity varied across cell types within each tissue condition (Figure 4A). CRC profiles showed broad diversity distributions across epithelial, myeloid, plasma-cell, and stromal compartments, while ANT tissue exhibited a related but distinct pattern. In healthy colon, several cellular compartments showed low or near-zero diversity, consistent with dominance by a limited number of detected viral annotations. Pairwise Mann–Whitney U tests with Benjamini–Hochberg correction identified significant differences between cell types within each condition (adjusted *p <* 0.05), as indicated by the compact-letter annotations in Figure 4A. These findings demonstrate that viral diversity varies across cellular compartments and tissue conditions rather than following a uniform distribution.

**Figure 4.**
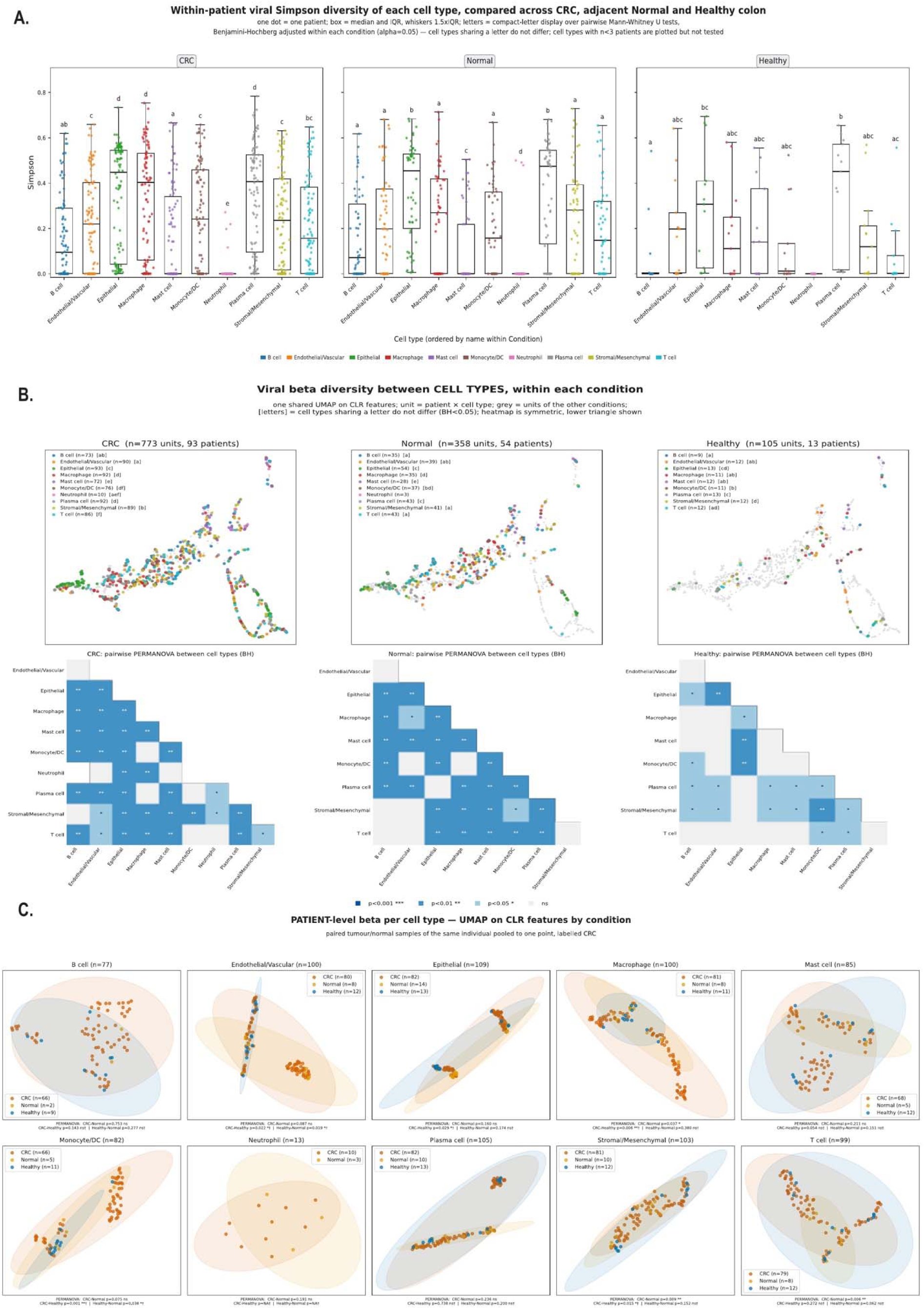
Participant-level viral alpha and beta diversity. (A) Simpson diversity by cell type within CRC, ANT, and healthy colon. Points represent participant-cell-type profiles; boxes show the median and interquartile range, whiskers extend to 1.5 times the interquartile range, and compact letters summarize BH-adjusted pairwise Mann–Whitney U tests within each condition. (B) UMAP visualization of CLR-transformed participant-cell-type profiles within each condition, with pairwise cell-type PERMANOVA performed on Aitchison distances. Heat maps show BH-adjusted significance categories. (C) Cell-type-specific UMAP views by tissue condition; matched CRC and ANT specimens from one participant are pooled into one point labeled CRC, as indicated in the artwork.

#### 3.2 Cell type structure in viral community composition

Beta-diversity analysis revealed cell-type-associated differences in viral-read composition within each tissue condition (Figure 4B). The participant-by-cell-type CLR profiles comprised 773 observations from 93 CRC participants, 358 from 54 ANT participants, and 105 from 13 healthy participants. Pairwise PERMANOVA with Benjamini–Hochberg correction identified multiple compositional differences among cell types, with more extensive compartmentalization in CRC and ANT tissues and a more limited but detectable pattern in healthy colon. These findings indicate that viral-read composition varies across cellular compartments rather than forming a uniform tissue-wide profile.

#### 3.3 Condition-associated compositional differences

Viral-read composition also differed among tissue conditions in a cell-type-dependent manner (Figure 4C). For visualization, matched CRC and ANT profiles were represented by a single participant point labeled CRC. Endothelial/vascular profiles differed between CRC and healthy colon (*p =* 0.022) and between ANT and healthy colon (*p =* 0.019), but not between CRC and ANT tissue (*p =* 0.087). Epithelial profiles differed between CRC and healthy colon (*p =* 0.029). Macrophage profiles showed differences between CRC and both ANT (*p =* 0.037) and healthy tissue (*p =* 0.006), while monocyte/dendritic profiles differed between CRC and healthy tissue (*p <* 0.001) and between ANT and healthy tissue (*p =* 0.038).

Additional differences were observed in stromal/mesenchymal profiles between CRC and ANT (p = 0.009) and CRC and healthy tissue (p = 0.015). T-cell profiles also differed between CRC and ANT tissue (p = 0.006). In contrast, B-cell, plasma-cell, and mast-cell profiles showed no significant compositional differences, while neutrophils were evaluated descriptively because healthy observations were unavailable. Together, these findings indicate that CRC-associated differences in viral-read composition vary across cellular compartments, with macrophage, stromal/mesenchymal, and T-cell profiles showing additional distinctions between tumour and ANT tissue.

### 4. Viral load and viral presence/absence feature sets support tissue-condition classification

#### 4.1 Viral load features classify tissue condition in repeated cross-validation

We evaluated whether cell-type-resolved viral-read-load profiles could distinguish CRC (94), ANT (54), and healthy (13) participant-condition profiles using 189 cell-type × virus features and 20 repetitions of stratified fivefold participant-grouped cross-validation (100 held-out test folds; Figure 5). The complete feature matrix is provided in Table S4 (viral-load sheet). Random forest achieved a mean macro-AUC of 0.957 ± 0.028 and accuracy of 0.840 ± 0.058, while CatBoost achieved a macro-AUC of 0.948 ± 0.031 and macro-F1 of 0.821 ± 0.093. Random forest one-versus-rest AUC values were 0.947 for CRC, 0.935 for ANT, and 0.990 for healthy tissue (Figure 5A). Distributions of the classification performance metrics are shown in Figure 5B.

**Figure 5.**
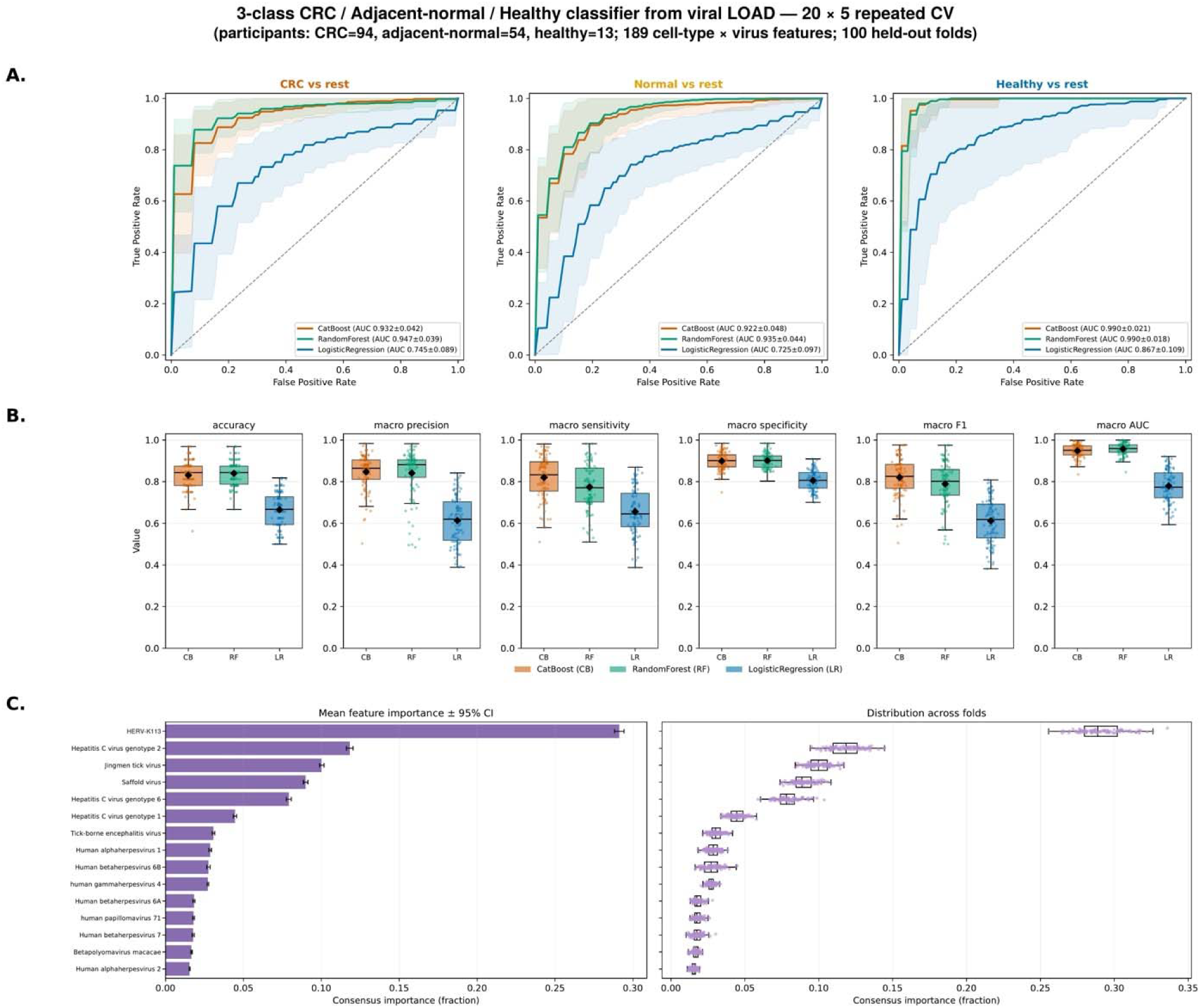
Repeated cross-validation of tissue-condition classification using viral-load features. (A) One-versus-rest ROC curves for CRC, ANT, and healthy colon from CatBoost, random forest, and logistic regression. (B) Distributions of accuracy, macro precision, macro sensitivity, macro specificity, macro-F1 and macro-AUC across 100 held-out test folds; black diamonds indicate means. (C) Global consensus viral feature importance across models.

Consensus feature importance ranked human endogenous retrovirus K113 (HERV-K113) highest, followed by hepatitis C virus genotype 2 (HCV-2), Jingmen tick virus (JMTV), Saffold virus, and hepatitis C virus genotype 6 (HCV-6) (Figure 5C). Class-specific permutation rankings are provided in Figure S6A. These results demonstrate that cell-type-resolved viral read load profiles contain substantial tissue condition information within the analyzed cohort, with several recurrent viral-reference features contributing most strongly to classification.

#### 4.2 Viral presence/absence features provide complementary classification performance

When the same 189 cell-type × virus features were represented as presence/absence rather than viral read load, all three classifiers retained substantial discrimination among tissue conditions (Figures S7A and S7B; Table S4, presence/absence sheet). Random forest achieved a mean macro-AUC of 0.931 ± 0.036 and accuracy of 0.824 ± 0.056, while CatBoost and logistic regression achieved macro-AUCs of 0.916 ± 0.042 and 0.905 ± 0.042, respectively. Logistic regression showed the highest mean macro-F1 (0.739 ± 0.116), closely followed by CatBoost (0.734 ± 0.111).

Global feature importance ranked hepatitis C virus genotype 2 (HCV-2) highest, followed by Saffold virus, HCV-6, HERV-K113, HCV-1, tick-borne encephalitis virus (TBEV), and Jingmen tick virus (JMTV) (Figure S7C). Class-specific importance rankings and complete performance distributions are provided in Figures S6B and S6C, respectively. Although random forest macro-AUC was somewhat lower than with quantitative viral-load features (0.931 versus 0.957), the binary representation retained considerable classification performance. Therefore, these results show that both the magnitude and occurrence of cell-type-resolved viral signals contain information for distinguishing tissue conditions within the analyzed cohort.

### 5. An external proteomic screen identifies annotation-compatible peptide matches

We examined an independent CRC proteomics cohort comprising 44 paired tumour and ANT acquisitions from 22 participants for peptide matches compatible with 20 candidate viral groups (Figure 6). All 20 candidates had at least one compatible peptide match in two or more acquisitions. EBV, HHV-6A, HHV-6B, CMV/HHV-5, HSV-1, and HSV-2 were represented in all 44 acquisitions; HHV-7, cowpox virus, and SuHV-1 in 43/44; norovirus GIII* in 42/44; HCoV-229E* and TBEV in 38/44; HCV* and Saffold virus in 37/44; HAdV-C* in 24/44; HIV-1 in 11/44; TTMV-9* in 8/44; rhinovirus A* in 7/44; and rhinovirus C* and astrovirus* in 2/44 each.

**Figure 6.**
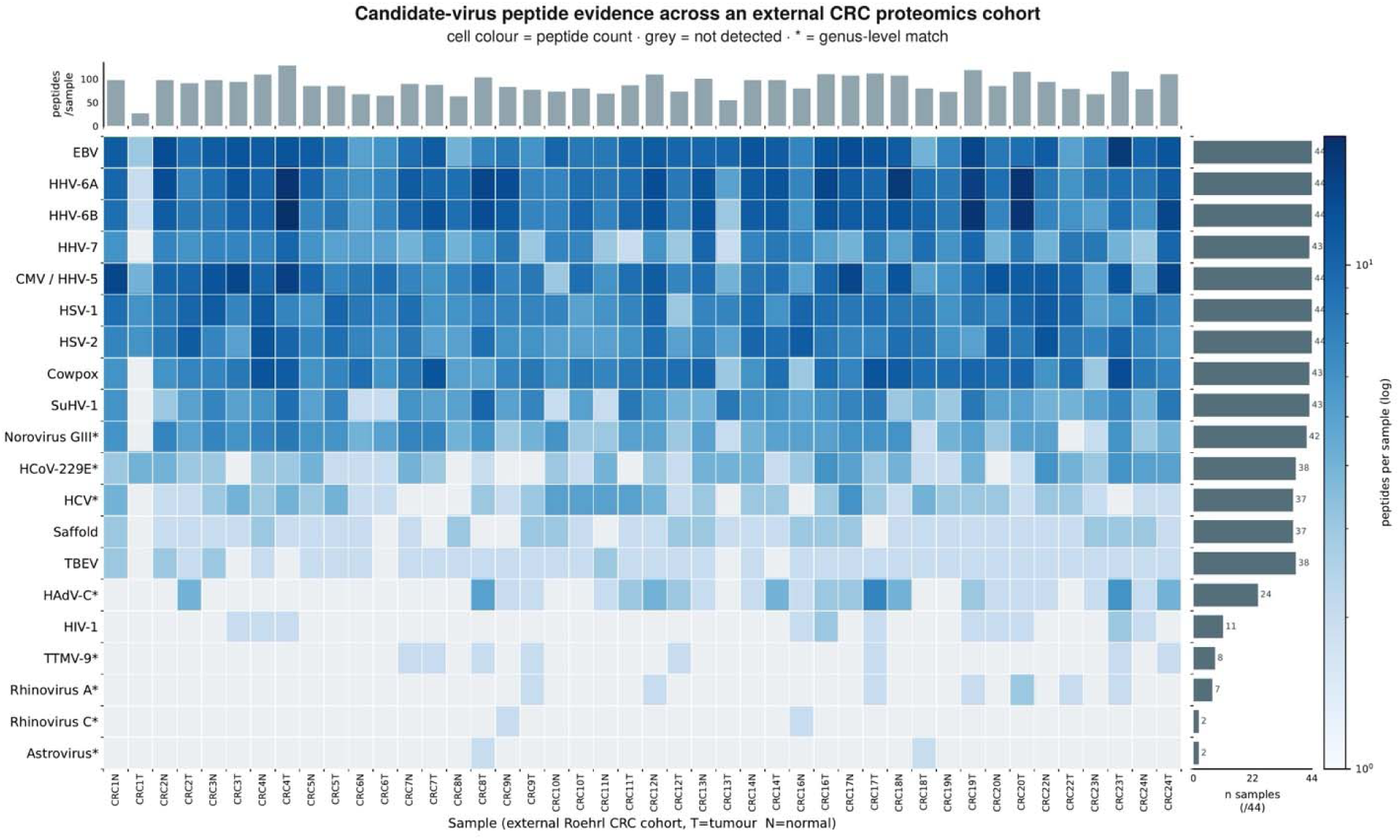
External proteomic peptide matches compatible with selected RNA-seq candidate annotations. The heat map shows accepted peptide-count signals for 20 candidate groups across 44 tumour and ANT acquisitions from 22 participants. Color denotes peptide count on a logarithmic scale, and gray denotes no detected peptide. An asterisk denotes a broader taxonomic match, including genus-level HCV.

Among these candidates, EBV and HCV also showed prominent viral patterns in the transcriptomic analysis. The independent proteomic screen therefore identified peptide matches compatible with several viral groups detected in the single-cell data. Asterisks indicate broader taxonomic assignments, including genus-level HCV, which do not establish genotype-specific identification.

### 6. Computational spatial inference shows cross-condition and CRC/ANT-associated projected patterns

To visualize cellular context for selected viral RNA-associated signals, we generated computational spatial projections for ten candidate annotations in a tissue-wide CRC view and in epithelial and plasma-cell contexts across CRC, ANT, and healthy tissue (Figure 7). Signal was visualized as arcsinh-transformed load [arcsinh(x/5)], with a separate color scale for each candidate. The maps therefore support within-candidate comparisons across conditions and cellular contexts, but do not permit comparisons of absolute load between viral annotations.

**Figure 7.**
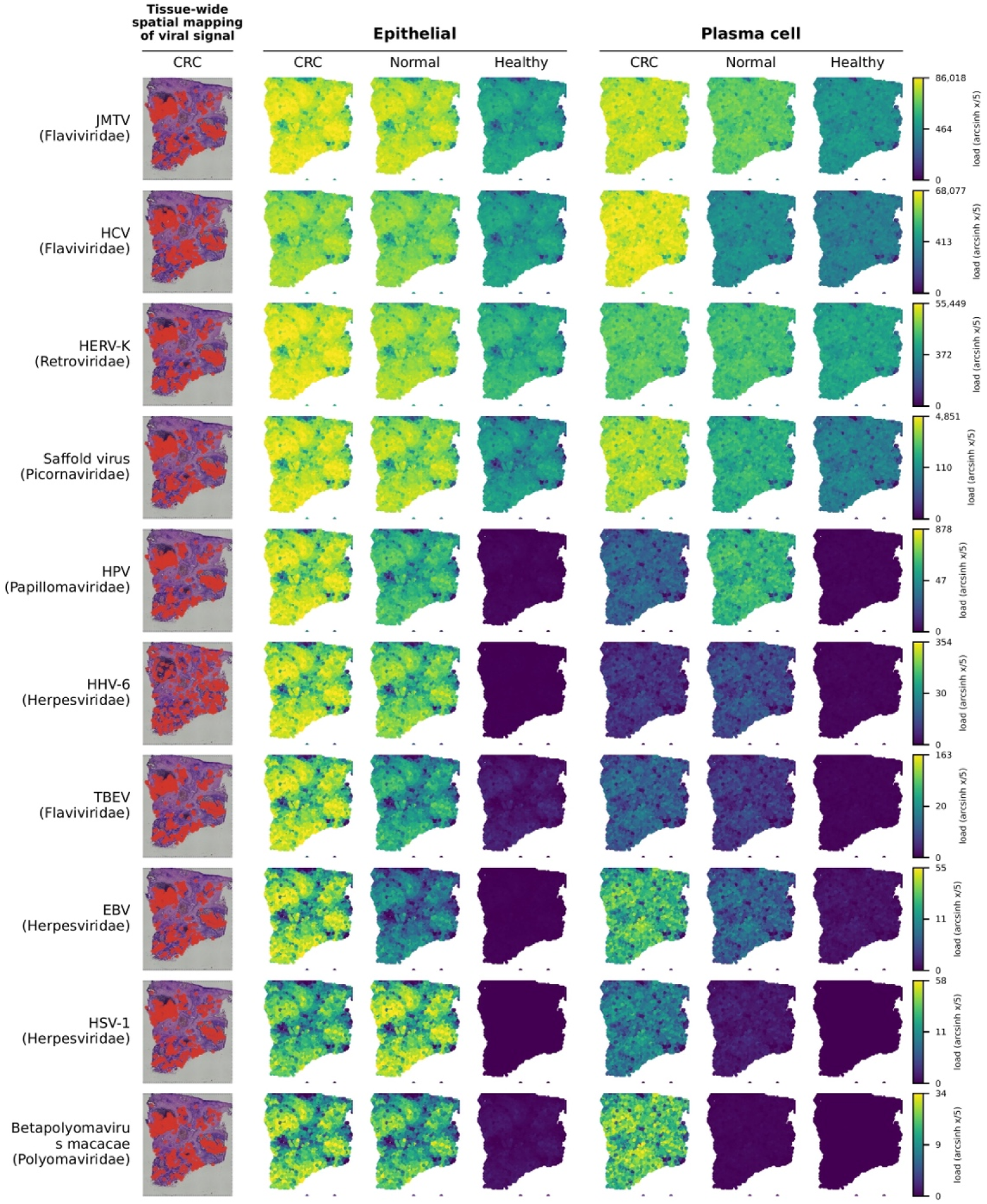
Computationally projected spatial patterns of selected viral RNA-associated signals in epithelial and plasma-cell contexts. Color encodes arcsinh-transformed load [arcsinh(x/5)] on a candidate-specific scale, so intensity is comparable only within a candidate row. These maps were generated using computational deconvolution and spatial inference.

#### 6.1 Candidates represented across all three tissue conditions

JMTV, HCV, HERV-K-associated signals, and Saffold virus showed detectable computationally projected viral-read loads in epithelial and plasma-cell contexts across CRC, ANT, and healthy tissue (Figure 7). Projected signals were generally less intense in healthy tissue than in CRC and ANT tissue, although they remained visible above background in several regions. These patterns were particularly widespread in epithelial projections, while JMTV, HCV, and HERV-K-associated signals also showed broad distributions in plasma cell projections.

#### 6.2 CRC/ANT-associated candidates

In contrast, HPV, HHV-6, TBEV, EBV, HSV-1, and Betapolyomavirus macacae showed predominantly CRC- and ANT-associated projected signals, with healthy tissue maps generally at or near background levels (Figure 7). These differences were most apparent in epithelial projections, suggesting distinct tissue condition patterns among the viral candidates.

### 7. Host transcriptional context, epithelial trajectory and cell–cell communication inferences

#### 7.1 Differential host-gene expression

Pooled differential-expression analysis of the atlas identified tissue-condition-associated differences in host-gene expression (Figure 8A), while cell-type-resolved analyses were performed separately within sequencing chemistries (Figure S8). In the pooled CRC versus ANT comparison, CLDN1, CXCL1, CXCL3, FOXQ1, ETV4, CDH3, and MMP7 showed higher expression in CRC, whereas BEST4, GUCA2A, GUCA2B, CA2, AQP8, and SLC26A2 were more highly expressed in ANT tissue. Cell-type-resolved differential expression results across the available tissue-condition comparisons are presented in Figure S8.

**Figure 8.**
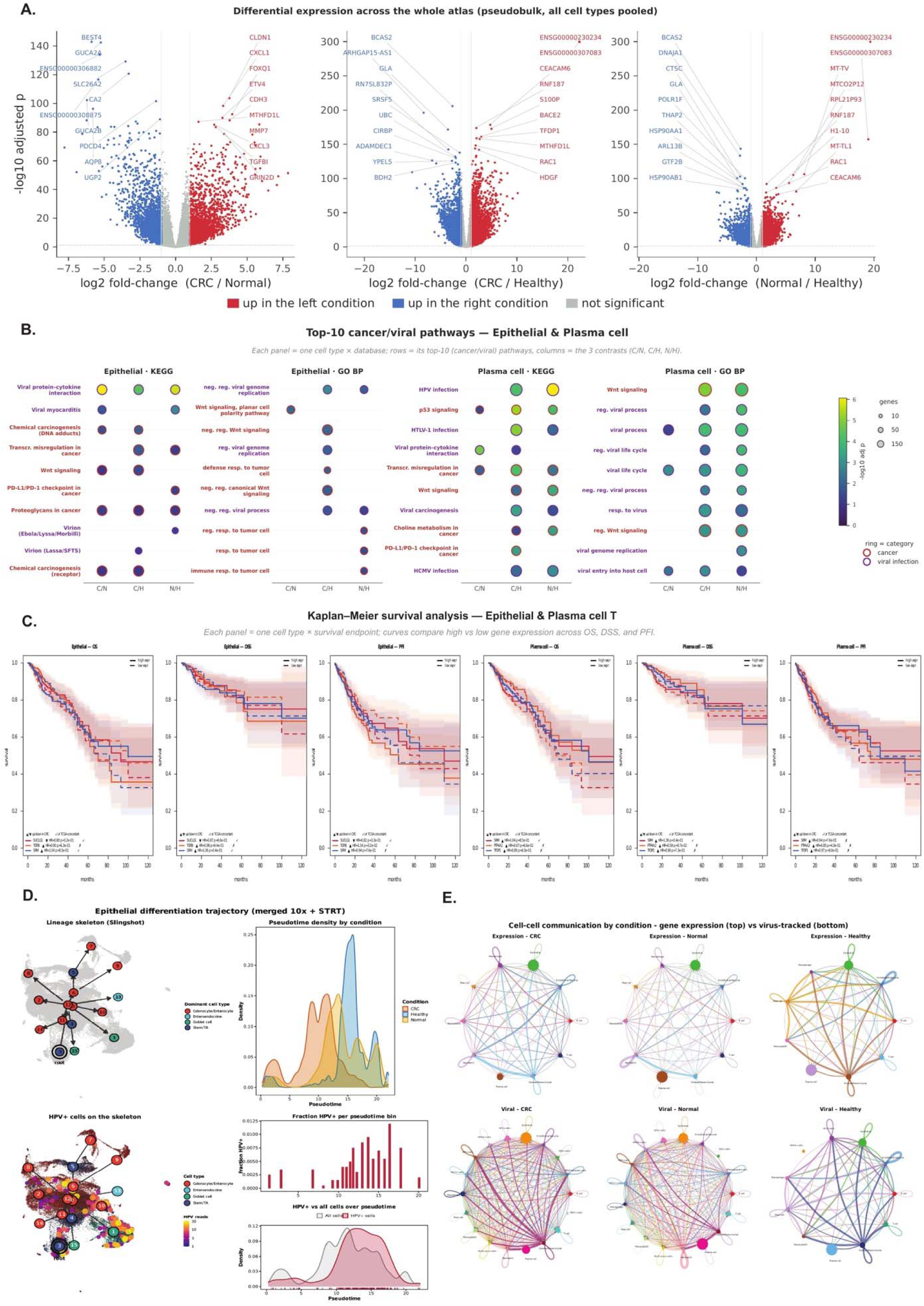
Host transcriptional, clinical-context, and inferred cell-state analyses of virus-associated RNA signals. (A) Whole-atlas pooled pseudobulk differential-expression volcano plots for CRC versus ANT (C/N), CRC versus healthy (C/H), and ANT versus healthy (N/H) tissue. The x-axis shows the log2 fold change for the stated left-versus-right contrast, and the y-axis shows the -log10 adjusted *p-value*. Red and blue points indicate higher expression in the left and right conditions, respectively; gray points are not significant. (B) Top ten cancer- and virus-category pathways in epithelial and plasma cells. Columns show C/N, C/H, and N/H contrasts; dot size represents the number of contributing genes, color represents the -log10 adjusted *p value*, and ring color identifies cancer or viral-infection category. Pathogen-named terms are curated host-gene annotations, not evidence of infection by the named virus. (C) Exploratory Kaplan-Meier curves for selected epithelial- and plasma-cell-derived host-gene candidates across overall survival, disease-specific survival, and progression-free interval in TCGA. (D) Inferred epithelial differentiation trajectory. Top, Slingshot lineage skeleton and pseudotime density by condition. Bottom, distribution of cells with HPV-associated reads on the skeleton, their fraction within pseudotime bins, and density relative to all epithelial cells. (E) CellChat-predicted condition-specific ligand-receptor network diagrams from host-gene expression (top row) and after inclusion of viral-read-defined cell groups (bottom row) for CRC, ANT, and healthy tissue.

#### 7.2 Cancer and virus category gene set enrichment

Host-gene enrichment analysis identified cancer and virus-related pathways in epithelial and plasma cell CRC versus healthy comparisons (Figure 8B). Epithelial cells showed enrichment of the KEGG pathway “Viral protein interaction with cytokine and cytokine receptor,” alongside Wnt signaling and cancer-related pathways. Plasma cells showed broader enrichment of p53, Wnt, HPV-labeled, HTLV-1-labeled, and other host response pathways. Complete enrichment results across the analyzed contrasts are provided in Figure S9 and Data S3.

#### 7.3 Clinical outcome association in TCGA-COADREAD

We evaluated the clinical-outcome associations of selected host genes using TCGA-COADREAD data (Figure 8C). The displayed candidates included SUCLG1, TGFBI, and SRM from epithelial cell analyses and SRM, PTP4A2, and TFDP1 from plasma cell analyses. Among these candidates, higher TGFBI expression was associated with a shorter progression-free interval (HR = 1.54, Cox *p =* 0.032). However, this association and the other nominal findings did not remain significant after correction for 81 survival tests. These results identify potential clinical associations that require further validation. Complete survival analyses are provided in Figure S10 and Data S4.

#### 7.4 Epithelial state trajectory of HPV-associated reads

To examine the distribution of HPV-associated reads across epithelial differentiation states, we reconstructed an inferred trajectory using merged 10x and STRT data (Figure 8D). The Slingshot trajectory originated from a stem/transit-amplifying (Stem/TA) associated cluster and branched toward colonocyte/enterocyte, goblet cell, and enteroendocrine states. Although HPV-associated cells were rare, they were distributed across multiple branches rather than confined to a single epithelial state. Their relative frequency was higher at intermediate to late pseudotime, peaking around pseudotime 16, while their density distribution was shifted toward later pseudotime compared with the overall epithelial cell population. These findings suggest that HPV reads are distributed across epithelial differentiation states, with greater representation at intermediate to late positions along the inferred trajectory.

#### 7.5 Inferred cell–cell communication

CellChat analysis revealed distinct predicted host ligand–receptor communication networks across CRC, ANT, and healthy tissues (Figure 8E). Incorporating viral-read-defined cell groups further revealed condition-dependent differences in network structure. CRC and ANT networks included HCV+, HHV-6+, HPV+, HSV-1+, EBV+, Betapolyomavirus macacae+, and multi-virus+ groups, with TBEV+ cells additionally represented in CRC. In contrast, healthy tissue contained fewer viral-read-associated groups (VRAS), with HCV+ cells participating in the predicted network and HHV-6+ cells showing no visible connections. Per-virus analyses also revealed distinct sender–receiver patterns (Figure S11). Together, these findings suggest that VRAS cell groups occupy different predicted communication contexts across tissue conditions, rather than sharing a uniform interaction pattern.

### 8. Segment-resolved profiling describes viral RNA-associated signals across colorectal anatomy

We summarized participant-level detection across the cecum, transverse colon, descending colon, and rectosigmoid, and placed the signals in broad cellular contexts (Figure S12A).

#### 8.1 Recurrent signals across colorectal segments

HCV-1, HCV-2, and Saffold virus showed consistently high detection frequencies across the four sampled colorectal segments (Figure S12A). HCV-2 was detected in 99–100% of participants across the cecum (n = 19), transverse colon (n = 19), descending colon (n = 7), and rectosigmoid (n = 70), while HCV-1 detection ranged from 91% to 100%. Saffold virus was detected in 61% of rectosigmoid participants and 100% of participants in the other three segments. These findings identify a recurrent group of viral reference-associated RNA signals across the sampled colorectal anatomy.

#### 8.2 Segment variable signals

Other viral candidates showed greater variation across colorectal segments (Figure S12A). JMTV was detected in all participants from the cecum and transverse colon, compared with 53% (37/70) in the rectosigmoid group. EBV and HPV-71 were detected in 79–84% of cecal and transverse colon participants, compared with 39% and 36%, respectively, in the rectosigmoid group. Similarly, TBEV detection ranged from 79% to 89% in the cecum and transverse colon, compared with 43% in the rectosigmoid. The seven descending colon participants also showed the recurrent viral signals described above. Together, these observations indicate greater regional variation in JMTV, EBV, HPV-71, and TBEV detection, with higher frequencies in the cecum and transverse colon than in the rectosigmoid. The circos plots mapped these VRAS signals across epithelial, immune, stromal/mesenchymal, and endothelial/vascular compartments (Figure S12A).

## Discussion

In this study, we recovered a cell-type-resolved layer of viral reference-associated RNA from human unmapped scRNA-seq reads while preserving host-transcriptomic information. Across 547 eligible libraries from 112 participants, including 528 libraries retained for viral analysis, this approach revealed distinct viral read patterns across colorectal cell types and tissue conditions. Building on Viral-Track and subsequent single-cell viral discovery methods ^3, 49, 50^, our contribution is a multi-cohort CRC framework that preserves barcode and UMI provenance and allows viral signals to be examined alongside host-cell states (Figure S12B).

Previous CRC virome studies reported bacteriophage-rich changes in fecal communities and multivariable signatures distinguishing CRC from controls 51-53. VLP-enriched mucosal studies instead emphasized eukaryotic viruses, including Anelloviridae and Torque teno virus 51. Bulk colorectal sequencing detected HERV-K113, HHV-6B, EBV, and HCV-associated sequences 13, while tumour RNA-seq identified low-level EBV signals without resolving their cellular origin 54. These approaches investigate different biological materials and cannot be directly equated. Unlike bulk profiling, our analysis links recoverable viral RNA to epithelial, immune, stromal, and vascular compartments, which can help distinguish tissue-wide associations from candidate cell-specific patterns.

HERV-K113 emerged as a prominent classifier feature and warrants particular attention. Unlike acquired infection, HERV-K113 is a germline retroviral insertion polymorphism: some individuals carry the provirus whereas others possess the empty insertion site ^14^. The frequent assignment of viral reads to HERV-K113, despite its known insertional polymorphism, raises an important question regarding locus-specific expression and differences between individuals carrying or lacking this provirus. This hypothesis has precedent: distinct HERV-K loci are differentially expressed in CRC ^55^, while demethylation-induced endogenous retroviral double-stranded RNA activates MDA5–MAVS–IRF7 signaling and suppresses CRC-initiating cells experimentally ^56^. Distinguishing HERV-K113-specific transcripts from homologous HERV-K sequences could reveal whether germline insertion status or transcriptional regulation is associated with CRC susceptibility, molecular subtype, or therapeutic response.

The epithelial HPV-71 and macrophage-associated HCV signals offer different mechanistic directions. High-risk HPV E6/E7 proteins disable p53/RB checkpoints in cervical cancer 57. Although HPV-71 has not been established as an oncogenic genotype or convincingly linked to cancer development, its epithelial-associated signal in our study raises the possibility of a previously unrecognized relationship with colorectal tumour biology. Examining HPV-71-specific transcription alongside p16, RB, and p53 activity could determine whether this association reflects a functional host–viral interaction. Experimental studies have linked HCV core protein to STAT3 activation, cyclin D1 expression, and WNT/β-catenin signaling in fibroblast and hepatoma cell models 58, 59. Macrophage-associated HCV-like reads therefore motivate testing inflammatory signaling and epithelial–myeloid crosstalk, not assuming colorectal HCV replication.

EBV signals in plasma cells versus epithelium are similarly consequential. EBV LMP1 activates EGFR–STAT3–ERK in nasopharyngeal carcinoma 60, yet colorectal in-situ studies have localized EBV predominantly to tumour-infiltrating lymphocytes rather than carcinoma cells 61. HHV-6 ORF-1 experimentally binds p53 and transforms fibroblasts 62, whereas HHV-7 infection of cultured peripheral blood mononuclear cells has been reported to increase TGF-β and TNF-α production, reduce IL-2 production, and inhibit lymphocyte proliferation 63. Thus, cell-specific herpesviral signals suggest separable tumor-cell and immune-regulatory mechanisms, including TGF-β-associated immune exclusion demonstrated in CRC models 64. EBER hybridization and viral-protein assays could directly resolve these alternative cellular mechanisms.

Additional candidates suggest focused immune assays: HSV-1 ICP47 blocks TAP-dependent MHC-I antigen presentation ^65^, and tick-borne encephalitis virus NS5 impairs interferon JAK– STAT signaling ^66^. Jingmen tick virus has been associated with neutrophilic inflammation in human infection ^67^, while Saffold-virus leader-protein research has examined interferon responses ^68^. Neither has an established CRC oncogenic role; sequence-specific confirmation remains essential.

Although these cell-associated viral signatures generate mechanistic hypotheses, their interpretation requires confirmation of taxonomic specificity and cellular localization, particularly for uncommon viral candidates and exploratory associations that did not withstand multiple-testing correction.

Clinically, these cell-resolved hypotheses could complement mismatch-repair/microsatellite-instability (MMR/MSI) stratification, which already guides PD-1 blockade in advanced CRC 69. Our internally discriminative viral classifiers warrant testing beyond source-study and sequencing-platform effects. Pairing candidate-positive cells with APC/WNT, TP53, KRAS/MAPK and antigen-presentation programs could prioritize mechanisms of tumour evolution and immune escape. Matched tumor–normal genotyping, viral RNA/DNA in-situ localization and independent cohort testing should determine whether candidate signatures track stage, molecular subtype, immune phenotype, recurrence or treatment response. Their value lies in identifying testable host–virus–cell interactions, rather than treating viral-read detection alone as proof of oncogenesis.

## Data and code availability

All datasets analyzed in this study are publicly available. scRNA-seq datasets and accession numbers are listed in the Key Resources Table and Table S2. The curated NCBI Virus RefSeq reference and curation provenance are provided in Data S1. Spatial analyses used the 10x Genomics Visium colorectal cancer dataset (Parent_Visium_Human_ColorectalCancer). Proteomic data are available through ProteomeXchange (PXD019504), and clinical-outcome analyses used TCGA-COADREAD data from the NCI Genomic Data Commons. All analysis code is available at: https://github.com/Mohammadamin-Mahmanzar/Repurposing_Unmapped_scRNA. Additional data are available at https://doi.org/10.6084/m9.figshare.33558034

## Ethics statement

This study used publicly available, de-identified datasets and did not involve new participant recruitment or direct interaction with human participants. No additional research ethics approval was required for this secondary analysis.

## Supplemental Information

Figure S1. Empirical selection and benchmarking of viral-read filtering thresholds

Figure S2. Synthetic-injection benchmarking of viral-read retention across the analytical workflow

Figure S3. Joint colon single-cell t-SNE atlas

Figure S4. Condition-stratified viral-read-compatible signals on the joint t-SNE atlas

Figure S5. Complete virus- and cell-type-specific viral-read profiles

Figure S6. Class-specific viral feature importance and repeated-cross-validation performance of viral-load and presence/absence classifiers

Figure S7. Repeated cross-validation of tissue-condition classification using viral presence/absence features

Figure S8. Cell-type-resolved differential expression across tissue conditions

Figure S9. Cell-type-resolved cancer- and virus-category host-pathway enrichment

Figure S10. Exploratory survival-expression analyses of selected CRC-associated host-gene candidates

Figure S11. Per-virus CellChat-predicted host ligand-receptor networks

Figure S12. Anatomical profile of viral RNA-associated signals and conceptual rationale for cell-resolved analysis

Table S1. Participant-level characteristics and sequencing-run contributions of the colorectal cohort

Table S2. Source-library metadata and protocol information for the colorectal single-cell RNA-sequencing datasets

Table S3. Software, reference resources and version information for the barcode-aware viral-read workflow

Table S4. Viral-load and viral presence/absence feature matrices used for tissue-condition classification. Provided as a single workbook containing two sheets: the viral-load feature matrix and the viral presence/absence feature matrix.

Data S1. Human-host viral reference sequences and curation provenance

Data S2. Virus–cell-type viral-load comparisons with raw and Benjamini–Hochberg-adjusted *p-value*s

Data S3. Complete reported pathway-enrichment statistics Data S4. Survival statistics for selected host-gene candidates

## Acknowledgements

We thank the investigators and participants of the ten source studies and acknowledge the public repositories and resources that supported this work, including NCBI SRA and GEO, NCBI Virus and RefSeq, Ensembl, ProteomeXchange/PRIDE and the NCI Genomic Data Commons.

We dedicate this study to the memory of David Baltimore (1938–2025), whose classification of viruses by genome type and replication strategy informed our viral-reference framework.

Artificial intelligence was used in two parts of this work. For cell-type annotation, four of the five consensus methods used machine learning or foundation models: CellTypist v1.7.1, scGPT v0.2.4, DeepCellSeek v1.0.0, and a large language model step using Claude Opus 4.8 through the mLLMCelltype v2.0.5. SingleR v1.10.0 was the fifth method. Each method contributed one vote to a five-method consensus label, and the authors reviewed every discordant call against canonical markers, as described in section 5 of the Methods. OpenAI ChatGPT assisted only with grammar, syntax, clarity, and readability. The writing tools were not used to generate data, perform statistical analyses, interpret results, or draw scientific conclusions. The authors reviewed and edited all resulting text and accept responsibility for the manuscript.

## Author Contributions

Conceptualization, M.M., K.R., K.K. and Y.D.; Methodology, M.M., K.R., K.K. and Y.D.; Software, M.M. and K.R.; Data Curation, M.M., K.R., K.K. and Y.D.; Formal Analysis, M.M., K.R., K.K. and Y.D.; Investigation, M.M., K.R., K.K. and Y.D.; Validation, M.M., K.R., K.K. and Y.D.; Visualization, M.M., K.R., K.K. and Y.D.; Writing – Original Draft, M.M., K.R., K.K. and Y.D.; Writing – Review & Editing, M.M., K.R., K.K. and Y.D.; Supervision, K.K. and Y.D.; Project Administration, K.K. and Y.D. All authors read and approved the final manuscript.

